# Profound synthetic lethality between SMARCAL1 and FANCM

**DOI:** 10.1101/2024.02.27.582393

**Authors:** Sumin Feng, Kaiwen Liu, Jinfeng Shang, Lisa Hoeg, William Yang, Sabrina Roy, Jordan T.F. Young, Wei Wu, Dongyi Xu, Daniel Durocher

## Abstract

DNA replication stress is a threat to genome integrity. The large SNF2-family of ATPases participates in preventing and mitigating DNA replication stress by employing their ATP-driven motor to remodel DNA or DNA-bound proteins. To understand the contribution of these ATPases in genome maintenance, we undertook CRISPR-based synthetic lethality screens with three SNF2-type ATPases: SMARCAL1, ZRANB3 and HLTF. Here we show that *SMARCAL1* displays a profound synthetic lethal interaction with *FANCM*, another ATP-dependent translocase involved in DNA replication and genome stability. Their combined loss causes severe genome instability that we link to chromosome breakage at loci enriched in simple repeats, which are known to challenge replication fork progression. Our findings illuminate a critical genetic buffering mechanism that provides an essential function for maintaining genome integrity.

## Introduction

The human genome encodes 33 SNF2-family ATP-dependent translocases^1, 2^. While this family is generally viewed as consisting of chromatin remodelling enzymes , at least 11 of the 33 SNF2-family proteins (CHD1L, RAD54B, RAD54L, RAD54L2, HLTF, SHPRH, ERCC6, ERCC6L/PICH, ERCC6L2, SMARCAL1 and ZRANB3) are directly implicated the maintenance of genome stability^3–15^, with the majority of the remaining 22 proteins also linked to various genome maintenance processes through their canonical chromatin remodelling function (e.g. CHD4, SMARCAD1 and ATRX)^16–18^. SNF2-family proteins are related to the Superfamily 2 (SF2) helicase superfamily and are composed of two conserved RecA-like ATPase lobes that lack a typical strand-separation wedge^19^, making them inactive as helicases. Instead, these proteins are united in their ability to use the energy derived from ATP hydrolysis to translocate on DNA^2, 20^. DNA translocation is at the core of the chromatin remodelling activities of many SNF2-family proteins. Exactly how this activity is employed in DNA repair and DNA replication fork management is poorly understood, although for some enzymes, such as ERCC6L, single molecule biophysical studies have revealed some clues^21^.

Given the sheer number of related enzymes, it is not surprising that the functional genomics of cancer cell lines and targeted genome-scale synthetic lethality screens have suggested the existence of genetic buffering among SNF2-family enzymes. For example, a highly penetrant synthetic lethal interaction has been found between *SMARCA2* and *SMARCA4*, which stimulated the development of SMARCA2-targeting agents for the treatment of *SMARCA4*-mutated cancers^22, 23^. It is likely that other such buffering interactions exist, hindering our ability to understand their biological function.

The SNF2 family members SMARCAL1, ZRANB3 and HLTF are all associated with the management of DNA replication forks and more specifically with fork reversal in response to DNA replication stress^3, 24^. They are recruited to stalled forks through interactions with other proteins, where they recognise and bind to specific DNA structures such as forked or single-stranded (ss) DNA^25–28^. All three can catalyze a phenomenon termed fork reversal, which is the process by which a replication fork, which can be viewed as a three-way junction, is remodelled into a four-way junction through the annealing of the nascent DNA^29^. Current models often lump these three translocases together as “fork remodellers”, but with some exceptions, their unique functions in genome maintenance remain elusive.

Since *ZRANB3*, *HLTF* and *SMARCAL1* are not essential in most human cell lines, we reasoned that we may be able to identify clues as to their unique functions by identifying genes whose mutation renders them essential for cellular viability. In other words, we sought to define their roles by charting their synthetic lethal interactions. To our surprise, we found that there is an absence of pairwise genetic buffering among these three genes but we discovered that loss of SMARCAL1 causes a profound dependence on an unrelated fork remodelling enzyme, FANCM. The converse is also true, with loss of SMARCAL1 causing lethality in *FANCM-* deficient cells. The lethality in cells lacking both SMARCAL1 and FANCM is due to rampant genome instability and chromosome breakage, particularly at A/T-rich regions or other genomic loci characterized by simple repeats. Our work is consistent with the possibility that SMARCAL1 and FANCM redundantly facilitate replication at a subset of genomic regions that are known to be problematic for the replisome.

## RESULTS

### SNF2-family fork remodeler synthetic lethality screens

To chart the genetic interaction network surrounding the genes encoding the main DNA replication fork remodelling SNF2 ATPases in human cells, we disrupted the *SMARCAL1*, *HLTF* and *ZRANB3* genes in RPE1-hTERT cells that constitutively express Flag-Cas9 and in which the p53 (*TP53*), and puromycin N-acetyltransferase (*PAC*) genes were also disrupted, yielding what will be referred to as the RPE1 *SMARCAL1^-/-^*, *HLTF^-/-^*and *ZRANB3^-/-^* cell lines, respectively (Figure S1A). We subjected these cell lines, and their parental counterparts (termed RPE1-hTERT *TP53^-/-^ PAC^KO^* Cas9 cells) to genome-scale CRISPR screens using the TKOv3 single-guide (sg) RNA library, and the screens were analyzed with BAGEL2^30^ and CRISPR Count Analysis (CCA) as previously described^31, 32^ to identify genes that are selectively essential in the mutated cell line (Figure 1A-D and Table S1).

**Figure 1.**
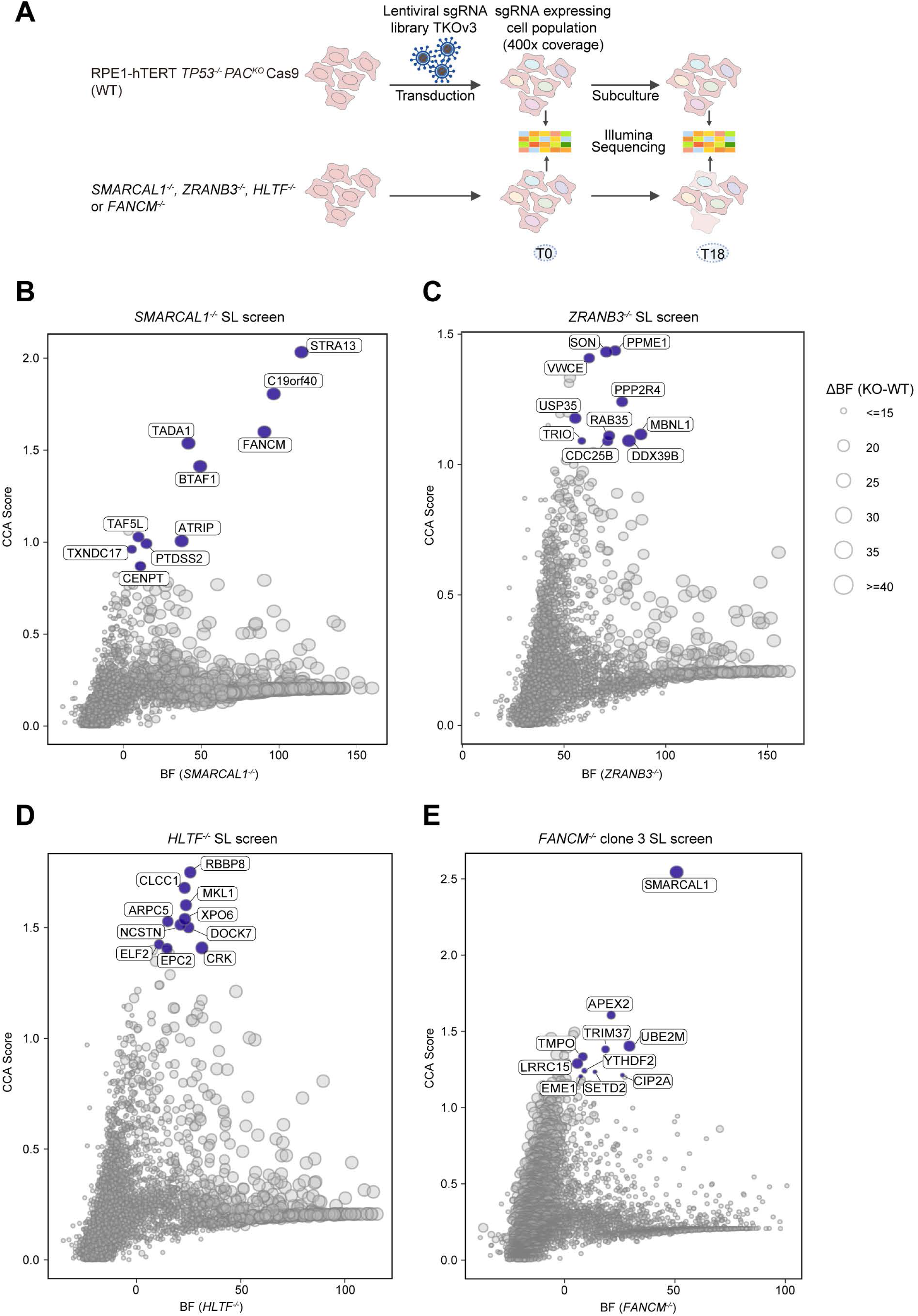
SNF2-family fork remodeler synthetic lethality screens. (A) Schematic of the isogenic, dropout CRISPR screens to identify synthetically lethal interactions with SMARCAL1, ZRANB3, HLTF and FANCM deficiency. WT, wild type. (B-E) Scatter plot of CRISPR Count Analysis (CCA) scores and Bayes factor (BF) values derived from BAGEL2 for the *SMARCAL1^-/-^* (B), *ZRANB3^-/-^*(C), *HLTF^-/-^* (D) and *FANCM^-/-^* clone 3 (E) isogenic synthetic lethal (SL) screens performed in RPE-hTERT *TP53^−/−^ PAC^KO^* Cas9 cells.

Synthetic lethal hits were defined as having a CCA score of above 1.0 and a Bayes Factor (BF) value of >10, which identified 30 (*HLTF*^-/-^ screen), 6 (*SMARCAL1^-/-^* screen) and 10 (*ZRANB3^-/-^*screen) genes. Analysis of gene ontology term enrichment using ShinyGO^33^ yielded DNA damage-associated terms for both the *HLTF^-/-^* (such as DNA double-strand break processing; GO:0000724) and *SMARCAL1^-/-^* screens (such as interstrand cross-link repair ; GO:0036297), whereas terms linked to cell cycle regulation and splicing (such as regulation of mRNA splicing, via spliceosome; GO:0048024 or cell division; GO:0051301) were enriched in the *ZRANB3^-/-^*screen (Figure S1B-D). Remarkably, three out of the six hits in the *SMARCAL1^-/-^* screen encoded either FANCM or its interacting partners FAAP24 (also known as C19orf40) and MHF2 (also known as STRA13) (Figure 1B). The other 3 hits in the *SMARCAL1^-/-^* screen were *ATRIP*, coding for the RPA-targeting subunit of the ATR kinase^34^, *TADA1* encoding a transcriptional co-activator subunit of the STAGA complex^35^ and, finally, the *BTAF1* gene, which codes for a SNF2 ATPase normally linked to transcriptional regulation^36^.

The striking genetic interaction between *SMARCAL1* and *FANCM, FAAP24 or MHF2* became the focus of this study. This was in large part due to the fact that like SMARCAL1, FANCM is also an ATP-driven fork remodelling enzyme, which suggested that these two enzymes may be acting redundantly at stalled or distressed replication forks.

Contemporaneously to the above screens, we also screened for synthetic lethal interactions with the fork remodelling enzyme *FANCM*. In this case, three independent *FANCM^-/-^* clones (#3, #12 and #15) were generated in RPE1-hTERT *TP53^-/-^ PAC^KO^* Cas9 cells, and their characterization is detailed in Figure S2A-C. The *FANCM^-/-^* screens were carried out as described above in clone #3 and #12. Using the same criteria to define hits, 19 (*FANCM^-/-^* clone #12) and 13 (*FANCM^-/-^* clone #3) genes were found to be synthetic lethal with *FANCM* for a total of 24 genes being a hit in one or the other screen. The GO enrichment terms for Biological Processes among the 24 hits retrieved many genome stability-related terms such as telomere maintenance (GO:0090737), DNA repair (GO:0006281) and DNA replication (GO:0006260) (Figure S2D). The two *FANCM^-/-^* screens had 8 hits in common (*SMARCAL1*, *APEX2*, *CIP2A*, *UBE2M*, *TRIM37*, *SETD2, NBN* and *POLG2*; Figure 1E, Figure S2E and Table S2). The *CIP2A-FANCM* genetic interaction was previously noted^31^ and was readily validated in two-color cell competition assays^37^ (Figure S3A) but by far the strongest genetic interaction detected in these screens was with *SMARCAL1* (Figure 1). Together, the screens converged on the possibility that *SMARCAL1* and *FANCM* genes display a strong and mutual genetic interaction.

### Validation of *SMARCAL1*-*FANCM* synthetic lethality

To validate the *SMARCAL1-FANCM* genetic interaction, we carried out clonogenic survival assays in which we inactivated FANCM, FAAP24 or MHF2 in parental or *SMARCAL1^-/-^* cell lines with two independent sgRNAs targeting each of their respective genes. We found that the loss of FANCM or FANCM-associated proteins selectively impaired the viability of SMARCAL1-deficient cells (Figure 2A,B). Similarly, depletion of SMARCAL1 (with two independent sgRNAs) in *FANCM^-/-^* cells also leads to pronounced lethality (Figure 2C). These results were corroborated in two-color cell competition assays (Figure S3B-D). The *SMARCAL1*-*FANCM* synthetic lethality was also confirmed in a second cellular background, the TERT-immortalized colon epithelial cell line, COL-hTERT *TP53^-/-^ PAC^KO^* Cas9 in which a *SMARCAL1*^-/-^ mutation was engineered (Figure 2D and Figure S3E,F). We conclude that *SMARCAL1* and *FANCM* display a robust and highly penetrant synthetic-lethal interaction in human cells.

**Figure 2.**
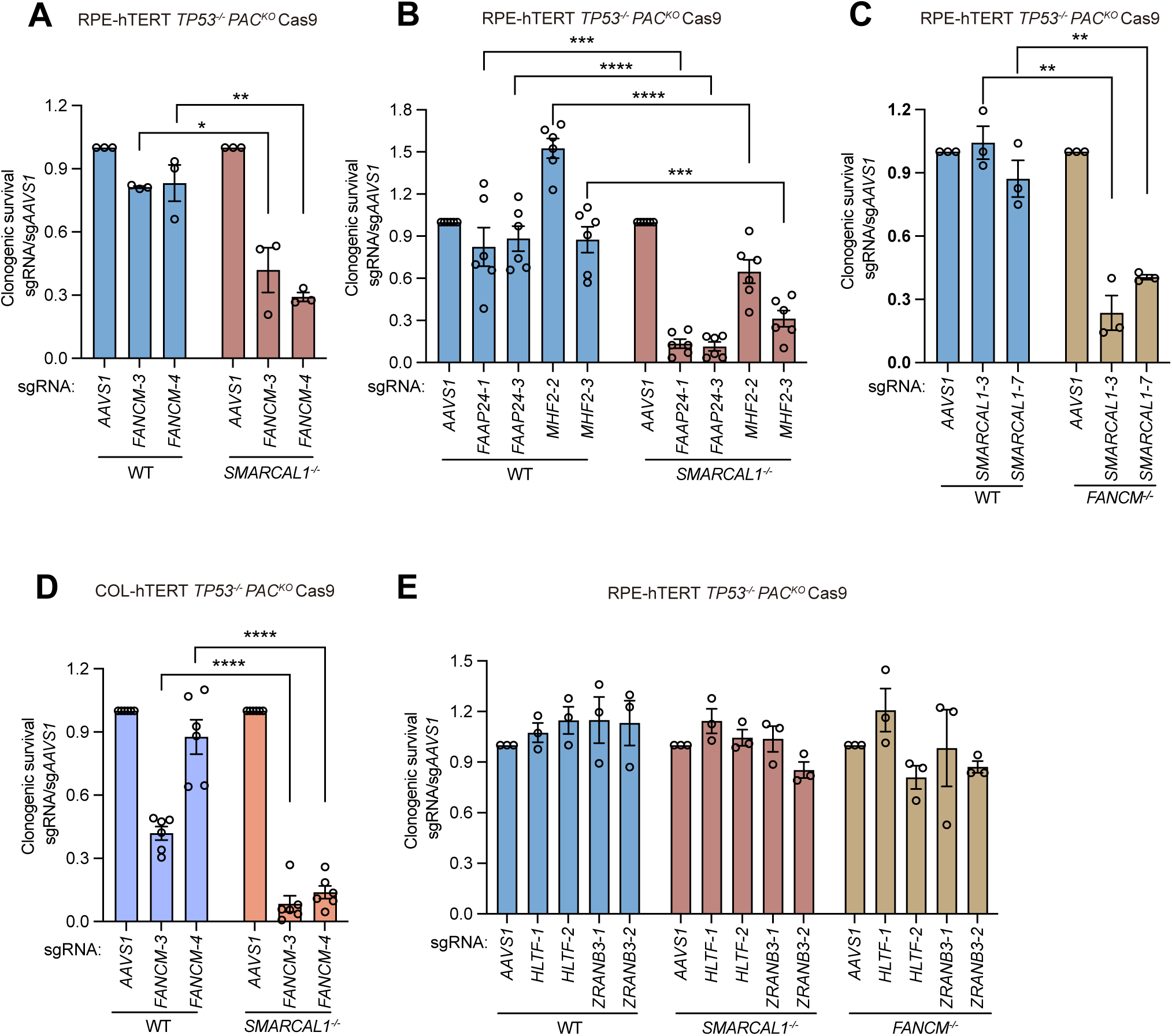
Characterization of the SMARCAL1 and FANCM-associated complex synthetic lethality. (A, B) Clonogenic survival of RPE-hTERT *TP53^-/-^ PAC^KO^* Cas9 parental (WT) and *SMARCAL1*^-/-^ cells upon transduction with a virus expressing an sgRNA targeting *AAVS1* (control), *FANCM* (A), *FAAP24* (B) or *MHF2* (B). (C) Clonogenic survival of RPE-hTERT *TP53^-/-^ PAC^KO^* Cas9 WT and *FANCM^-/-^* cells upon transduction with a virus expressing an sgRNA targeting *SMARCAL1* or *AAVS1* (control). (D) Clonogenic survival of COL-hTERT *TP53^-/-^ PAC^KO^* Cas9 WT and *SMARCAL1^-/-^* cells upon transduction with a virus expressing an sgRNA targeting *FANCM* or *AAVS1* (control). (E) Clonogenic survival of RPE-hTERT *TP53^-/-^ PAC^KO^* Cas9 WT, *SMARCAL1^-/-^* and *FANCM^-/-^* cells upon transduction with a virus expressing an sgRNA targeting *AAVS1* (control), *HLTF* or *ZRANB3*. For all panels, data is presented as the mean ± SEM (n=3) and was analyzed by a two-tailed t-test. *p <0.05, **p < 0.01, ***p < 0.005, ****p < 0.0001.

Since false-negatives are not uncommon in CRISPR screens, we also directly tested whether depletion of HLTF or ZRANB3 caused lethality in *SMARCAL1^-/-^*or *FANCM^-/-^* cells. We did not observe any impact of their disruption on the viability of either cell line, despite efficient gene editing (Figure 2E and Table S3). This indicates that *SMARCAL1* and *FANCM* have a unique genetic interaction among fork remodellers.

### Structure-function analysis of FANCM

In addition to its ATP-driven translocase domain, FANCM harbors multiple domains that engage in protein-protein interactions^38^ (Figure 3A). To assess which activity of FANCM supports the viability of SMARCAL1-deficient cells, we expressed FANCM variants in *FANCM^-/-^* cells from a lentiviral vector (Figure 3A). Expression of these FANCM variants was confirmed by immunoblotting (Figure 3B), with most expressing as well as the wild type FANCM control, with the exception of the variants lacking the MM3 and ERCC4 domains (ΔMM3 and ΔERCC4, respectively).

**Figure 3.**
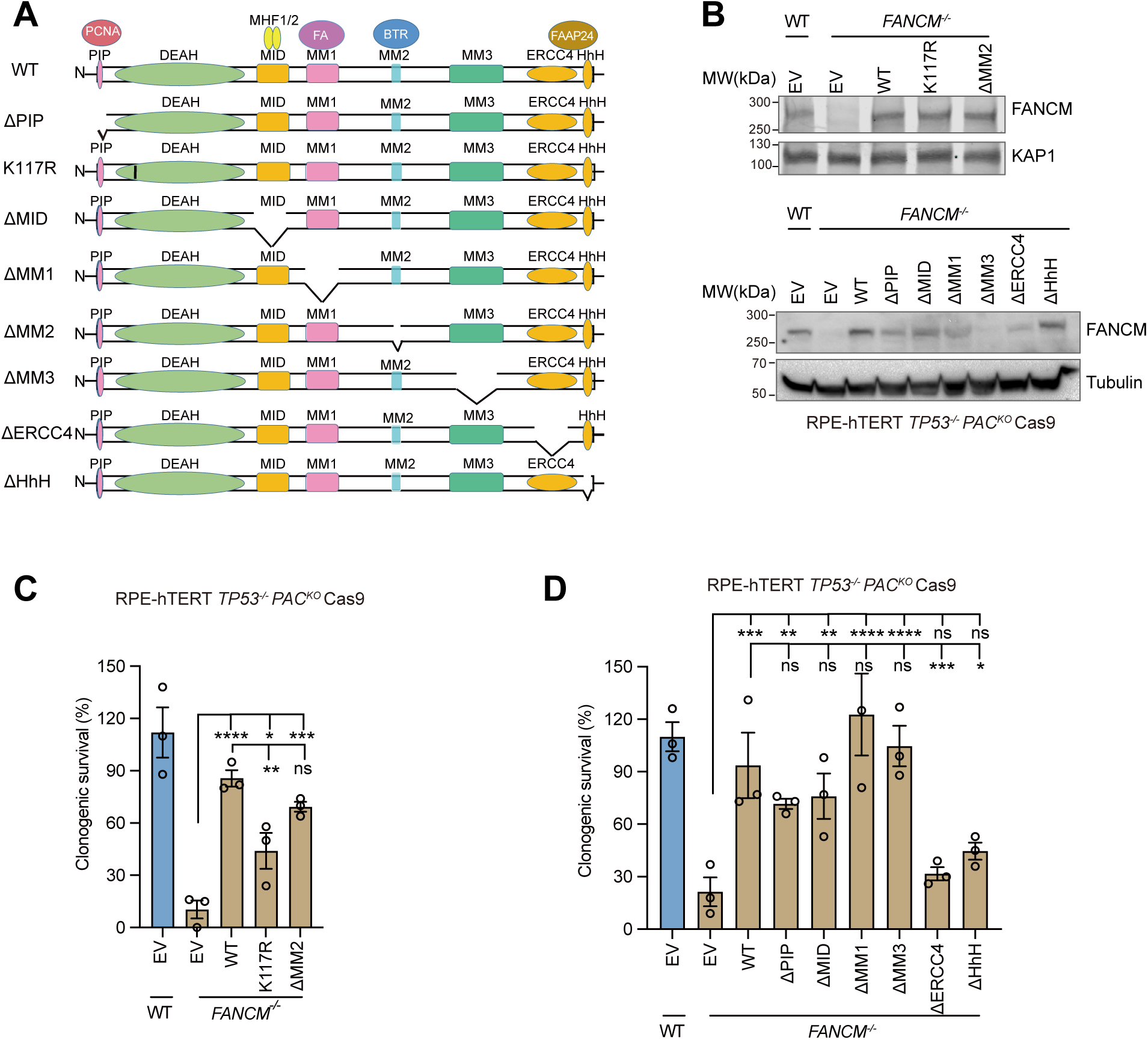
The ATPase activity of FANCM is required for cell viability in the absence of SMARCAL1. (A) Schematic of FANCM physical interactions, mutations and deletions. (B) Immunoblot analysis of whole-cell extracts of RPE1-hTERT *TP53^-/-^ PAC^KO^* Cas9 parental (WT) or *FANCM^-/-^* cells expressing wild type (WT) FANCM, the indicated mutants or an empty vector (EV). Lysates were probed for FANCM, KAP1 (loading control) and tubulin (loading control). (C) Clonogenic survival of RPE-hTERT *TP53^-/-^PAC^KO^* Cas9 parental (WT) or *FANCM^-/-^* cells with transgenes expressing an empty vector (EV), FANCM WT or its K117R and ΔMM2 mutants. Bars represent the survival of cells with a lentivirus-expressed SMARCAL1-targeting sgRNA relative to cells with an AAVS1-targeting sgRNA (control). Data is presented as the mean ± SEM (n=3) and analysis was performed with a one-way ANOVA. *p < 0.05, **p < 0.01, ***p < 0.005, ****p < 0.0001. ns, not significant. (D) As in (C) except cells were expressing an empty vector (EV), FANCM WT or its other deletion mutants (ΔPIP, ΔMID, ΔMM1, ΔMM3, ΔERCC4 and ΔHhH).

Complementation of *FANCM^-/-^* cells with lentivirally expressed wild-type (WT) FANCM renders the cells resistant to the lethality induced by SMARCAL1 depletion in both clonogenic or two-color competition assays (Figure 3C,D and Figure S4A-D). The impacts of mutation of the ATP translocase domain (K117R) or deletion of the ERCC4 (ΔERCC4) or HhH (ΔHhH) motifs were very clear and led to proteins that were largely defective in restoring viability to *FANCM^-/-^* cells depleted of SMARCAL1 (Figure 3C,D and Figure S4A-D). The ATPase-dead K117R mutant was not as defective as the empty-vector control condition in clonogenic survival assays, pointing to this variant retaining some activity. On the other hand, FANCM-ΔERCC4 and -ΔHhH were nearly completely inactive in both assays. The ERCC4 and HhH motifs form the heterodimerization surface that mediates the interaction of FANCM with FAAP24, which in turn promotes double-stranded (ds) DNA-binding and FANCM protein stability^39–41^. Since deletion of the HhH motifs generated a stably expressed FANCM protein (Figure 3B), we conclude that the FAAP24-FANCM interaction, and possibly dsDNA-binding by the FANCM-FAAP24 heterodimer, are necessary to support viability in the absence of FANCM.

Deletion of the PCNA-interacting PIP box (ΔPIP), the FA core complex-interacting MM1 domain (ΔMM1) and the MM3 domain (ΔMM3) all produced FANCM variants that promoted viability of *FANCM^-/-^* cells upon SMARCAL1 loss to levels that approached that of the wild-type protein (Figure 3D and Figure S4B,D). The situation was more nuanced with the FANCM-ΔMM2 and FANCM-ΔMID variants where expression of the former partially rescued the lethality caused by *SMARCAL1*-targeting sgRNAs (Figure 3C and Figure S4A,C) whereas for the latter, its expression largely rescued sensitivity to SMARCAL1 depletion in clonogenic assays (Figure 3D) but not in two-color competition assays (Figure S4B,D). While these results suggest that the interaction between FANCM and the BLM-TOP3α-RMI1/2 complex mediated by the MM2 domain^42^ is partially required to promote viability of SMARCAL1-deficient cells, depletion of BLM did not impact the FANCM-SMARCAL1 synthetic-lethal interaction (Figure S4E), pointing instead to the possibility that the ΔMM2 mutation may impact FANCM structural integrity. In contrast, despite the fact that the MHF1/2 proteins promote viability of *SMARCAL1^-/-^* cells, the FANCM-MHF1/2 interaction itself, mediated by the MID domain, may not be absolutely necessary for promoting the fitness of SMARCAL1-deficient cells.

### ATP-driven translocation by SMARCAL1 is essential in *FANCM^-/-^* cells

We next assessed which domains of SMARCAL1 promote viability of *FANCM*^-/-^ cells. We expressed, in RPE1 *SMARCAL1^-/-^* cells, Flag-tagged SMARCAL1 or variants with either a mutation in the ATP translocase domain (R764Q)^25^ or a deletion of the RPA-binding domain located in the N-terminal region (ΔN) (Figure 4A,B). We confirmed that unlike the wild-type and R764Q mutant, SMARCAL1-ΔN failed to colocalize with γ-H2AX following treatment with hydroxyurea (HU), as previously reported^25^ (Figure 4C,D). The resulting cell lines were then transduced with sgRNAs targeting *FANCM* (or *AAVS1* as negative control) and clonogenic survival was determined. We observed that expression of wild type SMARCAL1 or SMARCAL1-ΔN completely restored the viability of *SMARCAL1^-/-^* cells depleted of FANCM whereas SMARCAL1-R764Q was entirely ineffective in this assay (Figure 4E,F and Figure S4F). We conclude that the SMARCAL1 translocase activity promotes viability in the absence of FANCM independently of its interaction with RPA.

**Figure 4.**
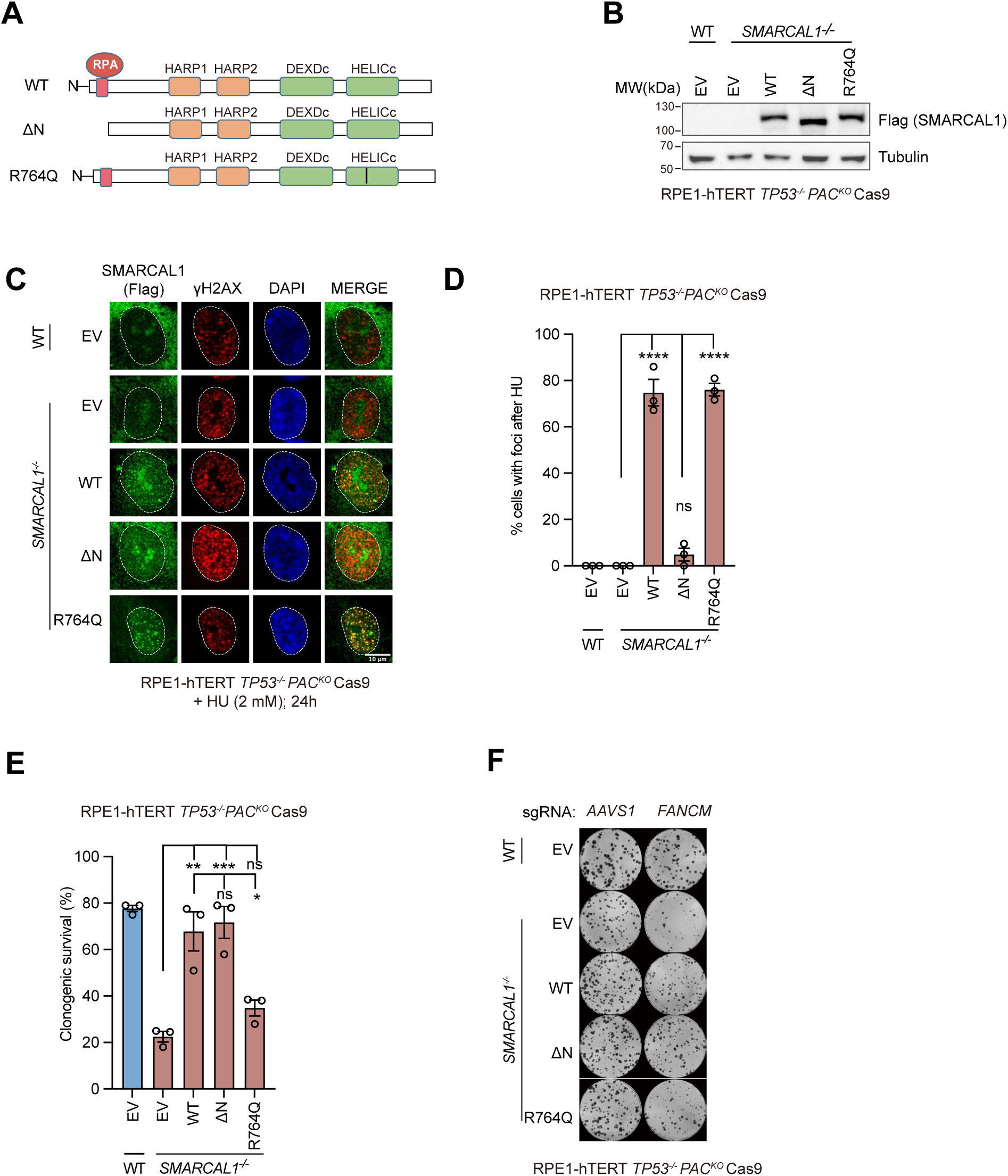
ATPase activity of SMARCAL1 is essential for cell viability in the absence of FANCM. (A) Schematic of SMARCAL1 physical interactions, mutations and deletions. (B) Immunoblot analysis of whole-cell extracts of RPE1-hTERT *TP53^-/-^PAC^KO^* Cas9 cells, parental (WT) or *SMARCAL1^-/-^*, expressing Flag-SMARCAL1 (WT), the indicated mutants or an empty vector (EV). Lysates were probed for Flag and tubulin (loading control). (C) Immunofluorescence analysis of HU-induced focus formation in RPE-hTERT *TP53^-/-^PAC^KO^* Cas9 WT or *SMARCAL1^-/-^* cells transfected with the indicated Flag-tagged SMARCAL1 variants. Cells were treated with 2 mM HU for 24 h, fixed and probed with antibodies to Flag and γH2AX and stained with DAPI. Representative micrographs are shown. Dashed lines indicate the nuclear boundaries determined by DAPI staining. Scale bar=10 μm. (D) Quantitation of (C). Data presented as the mean ± SEM (n=3) and was analyzed with a one-way ANOVA test. ****p < 0.0001,. ns, not significant. EV, empty vector. (E, F) Clonogenic survival of RPE-hTERT *TP53^-/-^ PAC^KO^* Cas9 WT or *SMARCAL1^-/-^* cells with transgenes expressing SMARCAL1 WT or mutants (ΔN and R764Q) and transduced with an sgRNA targeting *AAVS1* (control) or *FANCM*. EV, empty vector. (E) Quantitation of clonogenic survival. Data is presented as the mean ± SEM (n=3) and was analyzed with a one-way ANOVA test. *p < 0.05, **p < 0.005, ***p < 0.001,. ns, not significant. (F) Representative image of the experiment shown in (E).

### Rampant genomic instability in cells lacking both FANCM and SMARCAL1

To determine whether the root cause of the lethality observed following the combined loss of FANCM and SMARCAL1 was due to genomic instability, we first analyzed metaphase spreads of *FANCM^-/-^* cells in which we disrupted *SMARCAL1* via Cas9-mediated gene editing. We observed two striking phenotypes: first was a high level of chromatin breaks in the SMARCAL1/FANCM co-depleted cells (Figure 5A,B) and second, we found high levels of abnormal chromosome morphology, principally railroad-track chromosomes and chromosomes with premature chromatid separation, both suggestive of defective sister-chromatid cohesion (Figure 5C,D). As DNA replication stress is known to induce cohesion fatigue^43^, we suspect that both phenotypes originate from DNA replication-associated DNA lesions that are transmitted from interphase to mitosis. In support of this possibility, we observed accumulation of γ-H2AX-colocalizing CIP2A foci in mitotic cells lacking both FANCM and SMARCAL1 (Figure 5E,F). Indeed, CIP2A and TOPBP1 form a mitotic-specific complex that localizes at DNA lesions that are either transmitted from interphase or are generated in mitosis, including lesions originating from under-replicated DNA^31, 44^. We also observed a striking accumulation of cells in the G2/M phase of the cell cycle in the SMARCAL1/FANCM co-depleted cells, suggesting activation of checkpoint signaling (Figure 5G).

**Figure 5.**
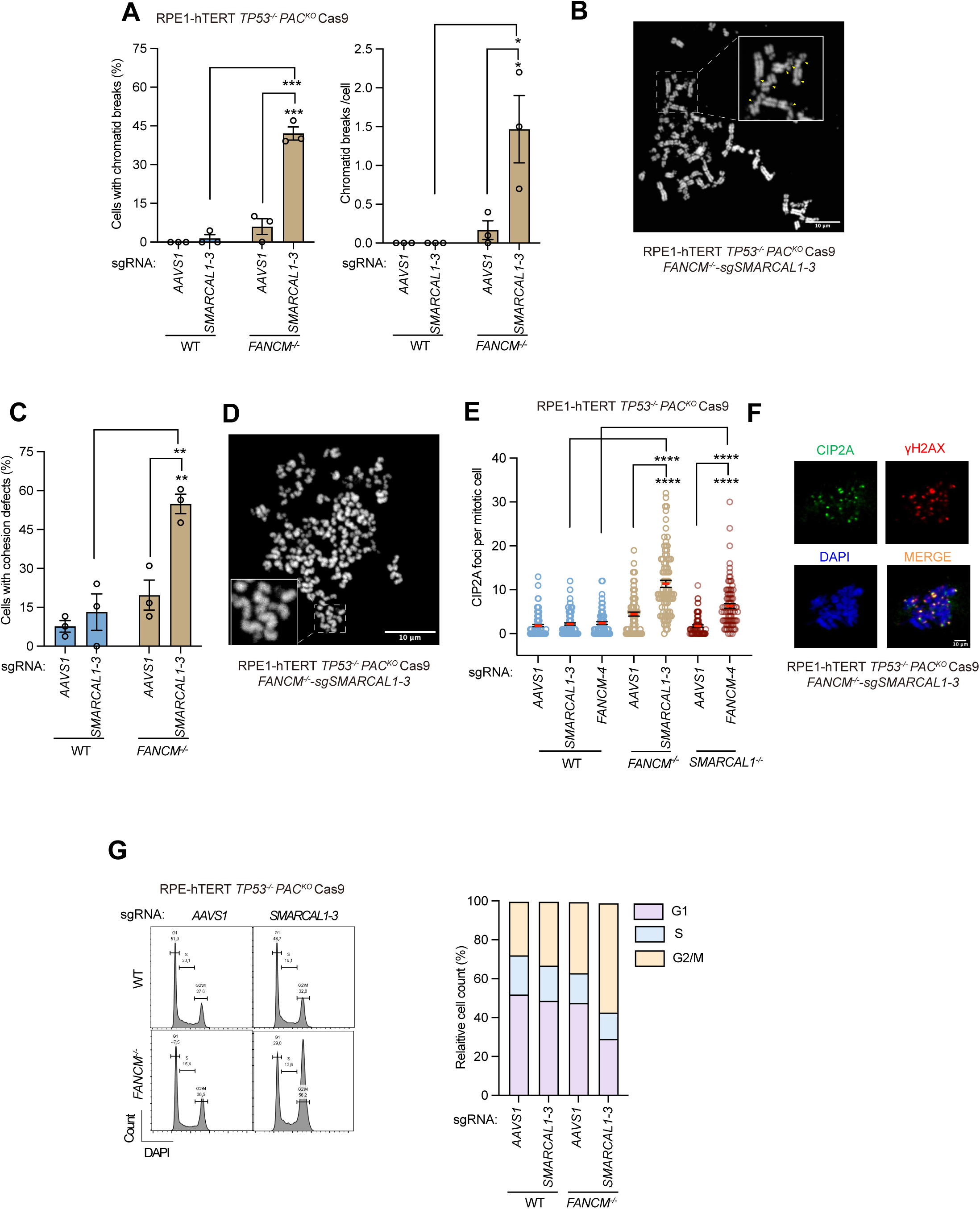
Combined loss of *SMARCAL1* and *FANCM* causes rampant DNA damage and genome instability. (A) Quantitation of chromatid breaks in metaphase spreads from RPE1-hTERT *TP53^-/-^PAC^KO^* Cas9 cells with the indicated genotypes upon transduction with a virus expressing an sgRNA targeting *SMARCAL1* or *AAVS1* (control). Results are shown as the percentage of cells with chromatid breaks (left) and number of chromatid breaks per cell (right). Data is represented as the mean ± SEM (n=3; 30 metaphases scored per experiment) and was analyzed with a two-tailed t-test. *p < 0.05, **p <0.01,. (B) Representative micrograph from the experiment quantitated in (A), showing a metaphase spread from *FANCM*^-/-^ cells transduced with a virus expressing an sgRNA targeting *SMARCAL1*. Inset highlights multiple broken chromosomes (yellow triangles). Scale bar = 10 μm. (C) As in (A) except the percentage of cells with cohesion defects was quantified. Data is presented as the mean ± SEM (n=3; 30 metaphases scored per experiment) and was analyzed with a two-tailed t-test. *p < 0.05, **p <0.01, ***p <0.001. (D) Representative micrograph from the experiment quantitated in (C), showing a metaphase spread from *FANCM*^-/-^ cells transduced with a virus expressing an sgRNA targeting *SMARCAL1*. Inset highlights premature chromatid separation. Scale bar = 10 μm. (E) Quantitation of CIP2A foci in mitotic RPE1-hTERT *TP53^-/-^PAC^KO^* Cas9 cells with the indicated genotypes upon transduction with a virus expressing an sgRNA targeting *SMARCAL1*, *FANCM* or *AAVS1.* Data is presented as the mean ± SEM (n=3) and was analyzed with a Mann–Whitney test, ****p < 0.0001. (F) Representative immunofluorescence micrographs from the experiment quantified in (E), showing a mitotic *FANCM*^-/-^ cell transduced with a virus expressing an sgRNA targeting *SMARCAL1* that has been probed with antibodies to CIP2A and γH2AX and stained with DAPI. The scale bar represents 10 μm. (G) Cell cycle distributions as determined by DAPI staining in RPE1-hTERT *TP53^-/-^PAC^KO^*Cas9 parental (WT) and *FANCM^-/-^* cell lines upon transduction with a virus expressing an sgRNA targeting *SMARCAL1* or *AAVS1*(control). Left, examples of FACS profiles. Right, percentage of cells in G1, S and G2/M phases from one representative experiment.

The fact that the loss either of SMARCAL1 or FANCM on their own causes some degree of DNA damage, made it difficult to assess DNA damage in interphase after depletion of SMARCAL1 in *FANCM^-/-^* cells (or vice-versa). To remedy this problem, we expressed SMARCAL1 fused to the conditional dTAG degron^45^ at its N-terminus in the parent RPE1-hTERT-derived *SMARCAL1^-/-^* cell line (yielding *dTAG-SMARCAL1* cells; Figure 6A). This system allowed us to knock out FANCM in *dTAG-SMARCAL1* cells and we ascertained that addition of dTAG-13 (0.5 μΜ) for 24 h caused near-complete degradation of SMARCAL1 in both the parental and *FANCM^-/-^ dTAG-SMARCAL1* cell lines (Figure 6Α). With this system, we observed an induction of DSBs, as detected by 53BP1 foci, in FANCM-deficient cells depleted of dTAG-SMARCAL1 (Figure 6Β,C). The increase in 53BP1 foci was dependent on cell cycling since it was not observed when SMARCAL1 was degraded in G1-synchronized *FANCM^-/-^* cells (Figure 6D,E). Together, these data suggest that the synthetic lethality between SMARCAL1 and FANCM is due to lethal induction of DNA damage during DNA replication, which is also supported by an observed increase in phosphorylated RPA (RPA pS4/pS8) (Figure 6F,G) as well as an accumulation of 53BP1 bodies in G1 phase in cells co-depleted of FANCM and SMARCAL1 (Figure 6H,I). Furthermore, *FANCM*^-/-^ cells depleted of SMARCAL1 are hypersensitive to hydroxyurea (HU), a DNA replication inhibitor, at times preceding their synthetic lethality (Figure 6J) further suggesting that DNA replication is at the root cause of the synthetic lethality.

**Figure 6.**
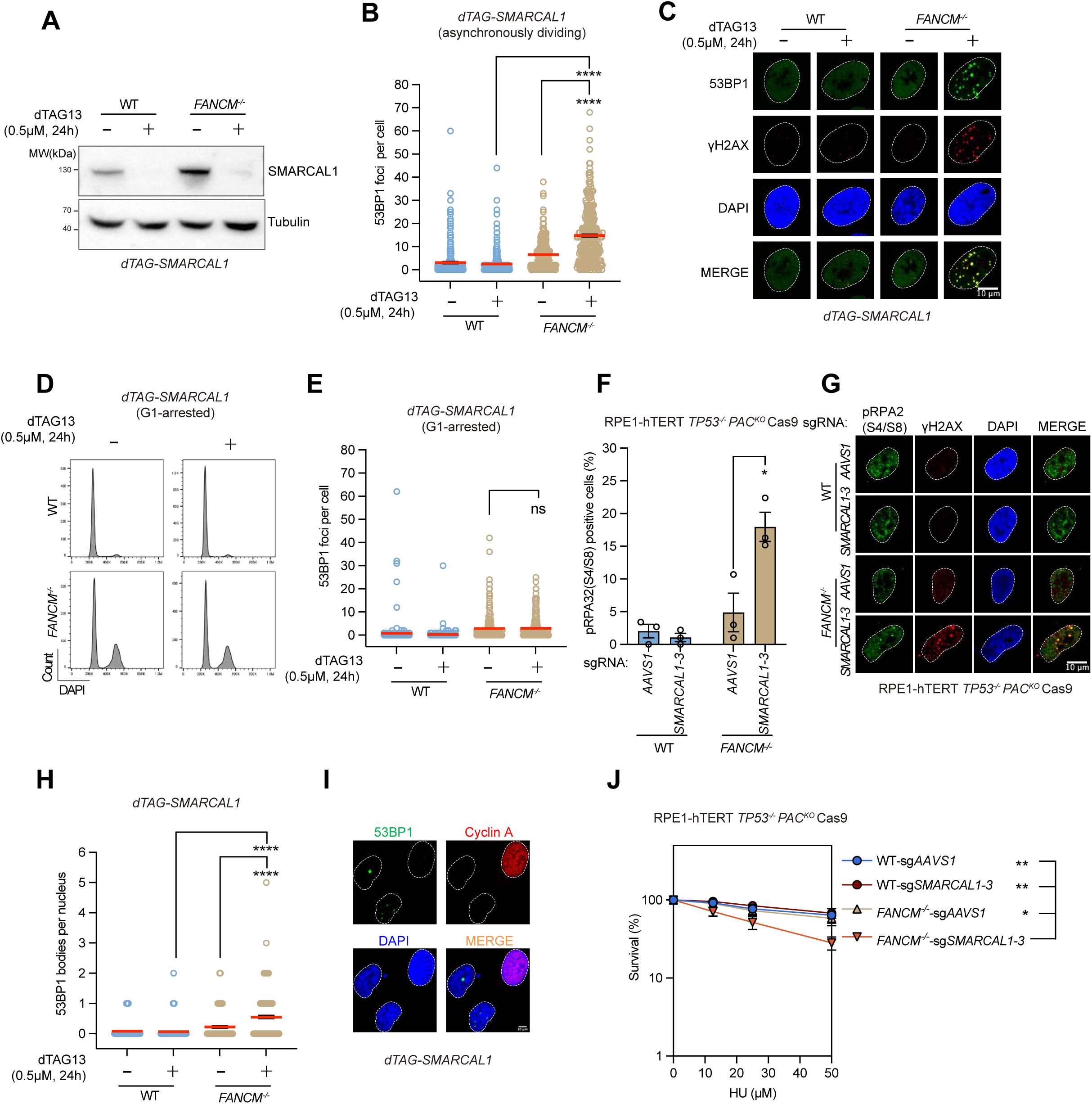
Combined loss of SMARCAL1 and FANCM causes DNA replication stress. (A) *dTAG-SMARCAL1* cells were created by transducing RPE1-hTERT-derived *SMARCAL1^-/-^* cells with a lentivirus expressing full length SMARCAL1 fused to the conditional dTAG degron at its N-terminus. FANCM was then knocked out in the parental (WT) *dTAG-SMARCAL1* cell line. Cells of the indicated genotype were treated with 0.5 μΜ dTAG-13 or DMSO (vehicle) for 24 h, and whole-cell extracts were analyzed by immunoblotting with antibodies to SMARCAL1 and tubulin (loading control). (B) Quantitation of 53BP1 foci in *dTAG-SMARCAL1* WT and *FANCM*^-/-^ cells with or without dTAG-13 treatment. Data is presented as the mean ± SEM (n=3) and was analyzed with a Mann–Whitney test, ****p < 0.0001. (C) Representative immunofluorescence micrographs of the experiment shown in (B). Cells were probed with antibodies to 53BP1 and γH2AX. The nuclear boundary was determined by DAPI staining (dashed line). Scale bar represents 10 μm. (D) FACS profiles of cell cycle distributions as determined by DAPI staining in *dTAG-SMARCAL1* WT and *FANCM^-/-^*cell lines with or without dTAG-13 treatment. All cells were synchronized in G1 phase by 1 μM palbociclib for 24 h and concurrently treated with either DMSO or dTAG13 (0.5 μM, 24 h). (E) Quantitation of 53BP1 foci in *dTAG-SMARCAL1* WT and *FANCM*^-/-^ cells treated as described in (D). Data is presented as the mean ± SEM (n=3) and was analyzed with a Mann-Whitney test. ns: not significant. (F) Quantitation of cells with pRPA32(S4/S8) foci in RPE1-hTERT *TP53^-/-^PAC^KO^* Cas9 parental (WT) and *FANCM^-/-^* cells upon transduction with a virus expressing an sgRNA targeting *SMARCAL1* or *AAVS1* (control). Data is presented as the mean ± SEM (n=3) and was analyzed with a two-tailed t test. *p < 0.05. (G) Representative immunofluorescence micrographs of the experiment shown in (F). Cells were probed with antibodies to phosphorylated RPA (pRPA32(S4/S8)) and γH2AX. The nuclear boundary was determined by DAPI staining (dashed line). Scale bar = 10 μm. (H) Quantitation of 53BP1 bodies in *dTAG-SMARCAL1* WT and *FANCM*^-/-^ cells with or without dTAG-13 treatment. Data is presented as the mean ± SEM (n=3) and was analyzed with a Mann– Whitney test, ****p < 0.0001. (I) Representative immunofluorescence micrographs of the experiment shown in (H). Cells were probed with antibodies to 53BP1 and cyclin A. The nuclear boundary was determined by DAPI staining (dashed line). Scale bar = 10 μm. (J) The survival curve of RPE1-hTERT *TP53^-/-^PAC^KO^* Cas9 parental (WT) and *FANCM^-/-^* cells upon transduction with a virus expressing an sgRNA targeting either *SMARCAL1* or *AAVS1*(control) and cultured under a gradient of HU treatments. Percent survival is displayed on a log scale. Data is presented as the mean ± SD (n=3) and was analyzed with a two-way ANOVA test, *p < 0.05, **p < 0.01.

### Mapping at-risk loci in FANCM/SMARCAL-deficient cells

To identify the loci prone to breakage when both FANCM and SMARCAL1 are absent, we carried out END-seq^46, 47^ in *dTAG-SMARCAL1* parental and *FANCM^-/-^* cell lines. These cells were treated with dTAG-13 for 24 h to deplete dTAG-SMARCAL1, or treated with DMSO as vehicle to maintain SMARCAL1 expression. After sequencing and peak-calling, the analysis confirmed that DMSO-treated *FANCM^-/-^*cells displayed a higher load of DSBs than dTAG13-treated parental cells, as determined by the number of peaks identified, and that dTAG-SMARCAL1 depletion in *FANCM^-/-^* cells greatly exacerbated the DSB load, as expected (Figure 7A). We identified 7360 peaks in the *dTAG-SMARCAL1 FANCM^-/-^*cell line when SMARCAL1 was depleted, and motif enrichment analysis showed striking enrichment for motifs such as poly(A/T) sequences, GG sequences and other simple repeats (Figure 7A-D). Interestingly, simple-repeat motifs are also enriched in the breaks mapped in *FANCM^-/-^* cells where SMARCAL1 expression is maintained but their breakage is greatly increased when SMARCAL1 is removed (Figure 7A). Since simple repeats are known as regions that promote replication fork stalling due to their propensity to adopt secondary structures^48–51^, these data suggest that the essential function of FANCM and SMARCAL1 resides in facilitating the replication of these structures (Figure 7C).

**Figure 7.**
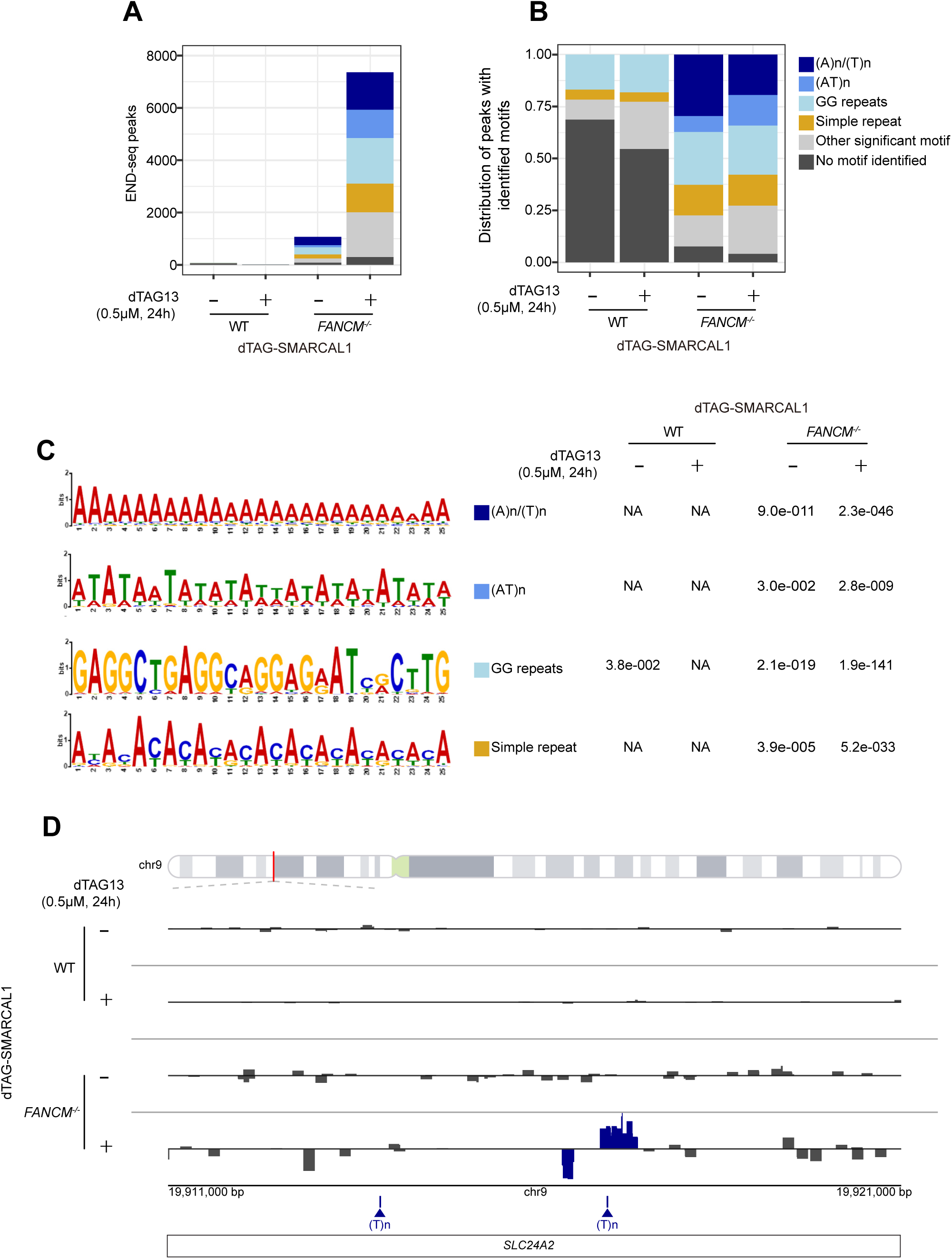
SMARCAL1/FANCM prevents breakage at simple repeat loci. (A) Quantitation and identification of END-seq peaks at (A)n/(T)n repeats (dark blue), (AT)n repeats (light blue), GG repeats (ice blue), simple repeats (yellow), other significant motif (grey, recurrent in multiple DSB regions) and DSBs with no identified motif (black) in *dTAG-SMARCAL1* WT and *FANCM*^-/-^ cells with or without dTAG-13 treatment (0.5 μM, 24 h). (B) The fraction of ends containing each motif in (A) at detected END-seq peaks in *dTAG-SMARCAL1* WT and *FANCM*^-/-^ cells with or without dTAG-13 treatment (0.5 μM, 24 h). (C) Representative motifs of analytic categories, as identified by performing STREME analysis on DNA sequences underlying the peaks identified in each cell line. P-value indicates the motif determined to be most representative of that analysis category from the STREME outputs. (D) Genome browser output of normalized read density for END-seq in *dTAG-SMARCAL1* WT and *FANCM*^-/-^ cells with or without dTAG-13 treatment (0.5 μM, 24 h). Shown are breaks at the *SLC24A2* locus on chromosome 9. Dark blue triangles represent the strand repeat annotation.

## Discussion

The DNA polymer can adopt various secondary structures, fold in higher-order chromatin arrangements and is the subject of multiple biochemical transactions, all of which can impede processive DNA replication. Cells overcome challenges to replication by expressing various factors that minimize the replication-blocking potential of DNA structures, remove problematic RNA-DNA hybrids, repair DNA damage and manage topological entanglements^52, 53^. One of the major challenges of the field has been to identify which proteins, among the multitude of replication fork accessory factors, are associated with a particular function. The work described here shows that SMARCAL1 and FANCM redundantly promote an essential function in the replication of simple repeat sequences such as poly(A/T) repeats. What underlies the imperiled nature of these simple repeats is unclear, as a subset of them is likely to be prone to DNA melting, whereas others (such as some G-rich sequences prone to breakage in FANCM/SMARCAL1 co-depleted cells) may instead be susceptible to hairpin formation or other secondary structures.

However, given that the essential activity imparted by FANCM and SMARCAL1 involves their ATP-dependent translocation, this crucial function most likely involves either remodeling of DNA structures or DNA-bound protein complexes. A clue as to what this activity is may be gleaned from considering the effect of SMARCAL1-ΔN, which is unable to bind to RPA but is fully able to suppress the SMARCAL1/FANCM synthetic lethality. Biochemical studies have shown that SMARCAL1 can use its ATP-driven motor activity to re-anneal RPA-coated bubbles in a manner that is independent of RPA-binding^25, 54, 55^. Given that a subset of breakage-prone simple repeats such as A/T rich sequences can form bubbles, one possibility may be that SMARCAL1 and FANCM use their ATP-driven motors to process bubble structures to prevent DNA replication fork stalling. An alternative possibility is based on the observation that the SMARCAL1-ΔN is also proficient at branch migration^55^, which is an activity shared by FANCM^56^. Indeed, given that branch migration is a component of fork reversal, the coupling of the “re-zipping” activity of SMARCAL1 and the branch-migration activity of SMARCAL1 or FANCM^56^ may allow forks to reverse at simple repeat-based obstacles to promote their resolution (or bypass). While this possibility is attractive, we note that the branch migration activity of FANCM is FAAP24-independent^56^, which is inconsistent with the observation that FAAP24, and the FAAP24-binding site on FANCM, are both essential in SMARCAL1-deficient cells. We expect that a more thorough comparative analysis of the ATP-driven activities of FANCM and SMARCAL1, guided by the structure-function studies presented here, may help uncover the exact type of structure(s) that are at risk of replication-associated breakage in FANCM/SMARCAL1 co-depleted cells. Such studies may also help explain why ZRANB3 or HLTF cannot compensate for the loss of these enzymes.

The profound synthetic lethality between FANCM and SMARCAL1 is also intriguing in the context of the association of both proteins with the alternative lengthening of telomeres, known as ALT^57^. SMARCAL1 minimizes DNA replication stress at ALT telomeres^58, 59^ and is essential in a subset of ALT cell lines such as SAOS-2, CAL72 and TM31 (depmap.org)^60^. Interestingly, this role of SMARCAL1 does not involve its RPA-interacting domain^58^ but involves its ATP-dependent translocase, suggesting that the role of SMARCAL1 in suppressing telomeric DNA replication stress may be instructive for understanding its role in *FANCM^-/-^*cells. Perhaps paradoxically, *SMARCAL1* mutations are also found in a subset of ALT tumors, such as in TERT promoter wild type glioblastoma^61, 62^, suggesting that under some contexts SMARCAL1 is a barrier to ALT initiation or maintenance. We expect that loss of FANCM will be particularly detrimental for *SMARCAL1*-mutated ALT tumors, and while this should be formally tested, we note that *FANCM* is essential in a wide range of *SMARCAL1* wild type ALT cell lines^63, 64^. However, in ALT, the essential function of FANCM depends on its ATPase activity and its interaction with the BTRR complex but is independent of FAAP24^63, 64^, arguing that this function may be unrelated to the role of FANCM in maintaining the viability of SMARCAL1-deficient cells. Nevertheless, we contend that understanding the molecular basis of the essential function mediated by the combined action of SMARCAL1 and FANCM may reveal unsuspected vulnerabilities to genome integrity and replisome progression, and that these insights may provide clues as to the role of these proteins in telomere maintenance.

## MATERIALS AND METHODS

### Cell culture

RPE1-hTERT *TP53^-/-^PAC^KO^* Flag-Cas9 ^37^, COL-hTERT *TP53^-/-^PAC^KO^* Flag-Cas9 ^65^ and 293T cells were grown in Dulbecco’s modified Eagle’s medium (DMEM; Gibco) supplemented with 10% fetal bovine serum (FBS; Wisent) and 1% Pen/Strep (Wisent). All cell lines were grown at 37°C with 5% CO_2_ and routinely tested negative for mycoplasma. The COL-hTERT *TP53^-/-^PAC^KO^* Flag-Cas9 *SMARCAL1^-/-^* and RPE1-hTERT *TP53^-/-^PAC^KO^* Flag-Cas9 *SMARCAL1^-/-^* cell lines were generated by transfection with *sgSMARCAL1* using Lipofectamine RNAiMAX (Invitrogen), followed by single clone isolation. Cell clones were validated by TIDE analysis^66^ and confirmed by immunoblotting. The RPE1-hTERT *p53^-/-^ PAC^KO^* Flag-Cas9 *FANCM^-/-^* cell line (constructed at Repare Therapeutics) were generated by transfection with *sgFANCM*, followed by single clone isolation. Cell clones were validated by TIDE analysis.

### Plasmids

For CRISPR/Cas9 gene editing, sgRNA targeting the desired gene was cloned into pLentiGuide-GFP-Puro, a modified form of Lentiguide-Puro (Addgene #52963), in which Cas9 was replaced with NLS-tagged GFP or mCherry by ligation into AgeI and BamHI sites. For SMARCAL1 complementation assays, the SMARCAL1 coding sequence was cloned into pHIV-NAT-T2A-hCD52 (a kind gift of R. Scully) by ligation into NheI and Xmal sites, then subcloned into pHIVT9-NAT-T2A-hCD52 by ligation into NotI and Xmal sites. pHIVT9-NAT-T2A-hCD52 was created by mutating the TATA box of the EF1a promoter (TATAA>CCCCC), which is reported to decrease the transcription of the strong wild type promoter^67^. The ΔN and R764Q mutants of SMARCAL1 were introduced by site-directed mutagenesis using Quikchange (Agilent). For FANCM complementation assays, pLVX-puro-3FLAG-FANCM-FL, K117R and ΔMM2 (Δ1219-1251) were gifts from Repare Therapeutics. All other mutants were introduced by site-directed mutagenesis using Quikchange. For *dTAG-SMARCAL1* cells, the SMARCAL1 coding sequence was cloned into pLEX_305-N-dTAG (Addgene #91797) by the Gateway system (Life Technologies/Thermo Fisher) according to the manufacturer’s protocol.

### Lentiviral transduction

Lentiviral particles were produced in 293T cells in 10-cm plates by co-transfection of 10 μg transfer vectors with packaging plasmids (5 μg pVSV-G, 3 μg pMDLg/RRE and 2.5 μg pRSV-REV) using calcium phosphate. Medium was refreshed 12–16 h later. Virus-containing supernatant was collected 36-48 h post-transfection, cleared through a 0.45 μm filter, supplemented with 8 μg/mL polybrene (Sigma) and used for infection of target cells.

### CRISPR screens

CRISPR screens were performed as described previously ^68^. RPE1-hTERT*p53^-/-^PAC^KO^*Flag-Cas9 parental and isogenic *FANCM^-/-^*, *SMARCAL1^-/-^*, *ZRANB3^-/-^* or *HLTF^-/-^* knock-out cells were transduced with the lentiviral TKOv3 library^69^ at a low MOI (∼0.3) and medium was refreshed 24 h later. Puromycin-containing medium (1.5 μg/ml) was added to select for transductants 48 h post-transduction. After an additional 48 h, corresponding to 4 d post-infection, cells were pooled together and divided into two sets of technical replicates, which defined the initial timepoint (T0). Cells were then subcultured every 3 d. Cell pellets were frozen at day 18 for gDNA isolation. Screens were performed in technical duplicates and library coverage of ≥ 400 cells per sgRNA was maintained at every step. gDNA from cell pellets was extracted using the QIAamp Blood Maxi Kit (QIAGEN) and genome-integrated sgRNA sequences were amplified by PCR using Q5 Mastermix (NEB Next UltraII, New Endland Biolabs M5044S). i5 and i7 multiplexing barcodes were introduced in a second round of PCR and final gel purified products were sequenced on Illumina NextSeq500 systems to determine sgRNA representation in T0 and T18 time points of each sample. To identify the list of genes whose inactivation causes loss of fitness, we compared sgRNA depletion using CCA^31^ and BAGEL2^30^ (T0 versus T18).

### Survival assays

For clonogenic survival assays, cells were seeded at a density of 500 or 750 cells/plate (depending on cell line and genotype) in 10-cm dishes and cultured as usual. Medium was refreshed after 7 d. After 14 d, medium was removed, cells were rinsed with PBS and then stained with 0.4% (w/v) crystal violet in 20% (v/v) methanol for 20 min. The stain solution was aspirated, and plates were rinsed twice with deionized water and air-dried. The colonies were counted using a GelCount instrument (Oxford Optronix), and data were plotted as surviving fractions relative to sgAAVS1-transduced controls. For HU sensitivity assays, cells were seeded in a 96-well plate using HU-free medium or medium containing the appropriate concentration of the drug. After 24 h, medium was removed, cand ells were cultured in HU-free medium. After 8 d, the medium in each well was refreshed and incubated for 30 min. 20 μL of CellTiter-Blue reagent (Promega) was then added to each well. The cells were shaken for 10 s and incubated for 3 h at 37°C. The fluorescence signal was recorded at 560/590 nm using an EnVision 2104 MultiLabel Reader (Perkin Elmer) and analyzed with the EnVision software.

### Two-color competitive growth assays

The two-color competitive growth assays were performed as previously described ^70^. Cells were transduced with lentivirus expressing NLS-mCherry-sgAAVS1 (control) or an NLS-GFP-sgRNA targeting *SMARCAL1* or *FANCM*. Medium was refreshed 24 h later. 48 h after transduction, virally transduced cells were selected with puromycin at 1.5 ug/mL for an additional 48 h. mCherry-and GFP-expressing cells were mixed in 1:1 (1000 cells each) and seeded in a 12-well plate. Cells were imaged for GFP and mCherry signals 24 h after initial plating (T=0d) and at the indicated time points (T=4d, 8d, 12d, 16d, 20d) using a 4X objective (InCell 6000 Analyzer, GE Healthcare Life Sciences). Segmentation and counting of GFP- and mCherry-positive cells were performed using an Acapella script (PerkinElmer).

### Antibodies

The following primary antibodies were used in this study at the indicated dilutions for immunoblotting (IB), immunofluorescence (IF), or fluorescence-activated cell sorting (FACS): rabbit anti-SMARCAL1 (1:1000 for IB; Abcam, ab154226), rabbit anti-ZRANB3 (1:1000 for IB; Proteintech, 23111-1-AP), rabbit anti-HLTF (1:3000 for IB; Abcam, ab17984), mouse anti-α-tubulin (1:1000 for IB; Calbiochem, CP06), rabbit anti-KAP1 (1:5000 for IB; Bethyl, A300-274A), goat anti-GFP (1:5000 for IF; a kind gift from Laurence Pelletier), mouse anti-γH2AX(S139) (1:2500 for IF; Millipore, 05-636), rabbit anti-γH2AX(S139) (1:500 for IF; Cell Signaling, 2577L), mouse anti-CIP2A (1:500 for IF; Santa Cruz, sc80659), rat anti-FLAG (1:1000 for IF; BioLegend, 637301), rabbit anti-53BP1 (1:2000 for IF; BD, A300-272A), mouse anti-Cyclin A (1:200 for IF; Santa Cruz, sc271682). Secondary antibodies for immunoblotting (IB) or immunofluorescence (IF) used in this study were as follows. For IB, IRDye 800CW goat anti-mouse IgG (1:5000; LiCOR, 926-32210), IRDye 680RD goat anti-rabbit (1:5000; LiCOR, 926-68071), HRP-conjugated sheep anti-mouse IgG (1:5000; GE Healthcare, NA931), HRP-conjugated goat anti-rabbit IgG (1:5000; Cedarlane, 111-035-144). For IF, AlexaFluor 488-goat anti-rabbit IgG (1:1000; Thermo Fisher Scientific, A11034), AlexaFluor 488-goat anti-mouse IgG (1:1000; Thermo Fisher Scientific, A11029), AlexaFluor 555-goat anti-mouse IgG (1:1000; Thermo Fisher Scientific, A21424), AlexaFluor 647-goat anti-mouse IgG (1:1000; Thermo Fisher Scientific, A21236), AlexaFluor 647-goat anti-rabbit IgG (1:1000; Thermo Fisher Scientific, A21244).

### Immunofluorescence

Cells were grown on coverslips for 24 h, and then exposed to the specified treatments or kept without any intervention. Cells were rinsed with PBS, and fixed with 4% paraformaldehyde (Thermo Fisher) for 10 min at room temperature. Cells were washed three times with PBS and subsequently permeabilized with 0.5% TritonX-100 in CSK buffer (100 mM NaCl, 20 mM HEPES pH 7.0, 3 mM MgCl_2_, and 300 mM sucrose) for 10 min at 4 °C. Cells were then rinsed three times with PBS, blocked with 5% BSA for 20 min, and incubated with primary antibodies for 90 min at room temperature. After washing 3 times with PBS, cells were stained with secondary antibodies for 45 min at room temperature and washed three times with PBS, mounted with DAPI-containing ProLong Gold mounting reagent (Invitrogen), and imaged using a Zeiss LSM780 laser scanning microscope with a 60X objective. Foci were manually counted.

### Cytogenetic analysis

Metaphase spreads were obtained from cells exposed to 100 ng/ml of KaryoMAX colcemid (Gibco/Thermo Fisher Scientific) for 2.5 h and harvested. Cells were then washed with PBS and resuspended in 75 mM KCl for 30 min at 37°C. After centrifugation, the supernatant was removed and cells were fixed by drop-wise addition of 1 ml of fixative (ice-cold methanol:acetic acid, 3:1). An additional 9 ml of fixative was then added, and cells were incubated at 4 °C for at least 16 h. Once fixed, metaphases were dropped onto glass slides and air-dried overnight, protected from light. To visualize chromosomal aberrations, slides were dehydrated in a 70%, 95% and 100% ethanol series (5 min each), air-dried and mounted in DAPI-containing ProLong Gold mounting reagent (Invitrogen). Images were captured on a Zeiss LSM780 laser-scanning confocal microscope.

### Cell cycle analysis (FACS)

Cells were transduced with a virus expressing an sgRNA targeting *SMARCAL1* or *AAVS1* (control). Cells were plated on 6-cm dishes to grow for 48 h before harvesting. The culture supernatant was collected, and cells were rinsed with PBS, trypsinized, and pooled with their respective culture supernatant. Cells were spun down, and pellets were washed twice with cold PBS and then incubated with analysis buffer (0.5 µg/ml DAPI in 1%BSA/PBS, 250 µg/ml RNaseA) for 30 min at room temperature in the dark. Cells were analyzed on an Attune NxT/CytKick Max autosampler (Thermo Fisher). Flow cytometry data were analyzed using FlowJo software. Live cells and subsequently single cells were selected and gated into G1, S, G2/M populations. To conduct the *dTAG-SMARCAL1* cell cycle analysis, WT and *FANCM^-/-^* cells lines were synchronized at G1 phase using 1 μM palbociclib for 24 h. Simultaneously, the cells were treated with either DMSO or 0.5 μM dTAG13 for the same duration.

### END-seq

END-seq experiments was performed essentially as previously described^47^ with the modifications detailed below. Briefly, 5 million cells were collected. DT40 cells were treated with 10 μM etoposide for 2 h, and collected as spike-in cells. RPE1 and DT40 cells were mixed at 50:1 and embedded in 0.75% low melting agarose plug. Afterwards, the polymerized agarose was treated with proteinase K (1 h at 50 °C, followed by 7 h at 37 °C), rinsed 3 times with plug wash buffer (10 mM Tris-Cl, pH 8.0, 50 mM EDTA, diluted in nuclease free water) and twice with TE buffer, and then subjected to RNAse A treatment for 1 h at 37 °C. Plugs were then washed three times with plug wash buffer, and four times with elution buffer (10 mM Tris-Cl, pH 8.0, in nuclease-free water). Next, the DSB ends were blunted, subjected to A-tailing and finally ligated with biotinylated hairpin adaptor 1. Following ligation with adaptor 1, DNA was recovered by melting the agarose plugs and then digesting the agarose with β-agarase enzyme (NEB, M0392L), using the manufacturer-recommended conditions. The extracted DNA was sheared to a length of ∼175 bp by sonication (ME220, Covaris) and biotinylated DNA fragments were purified using streptavidin beads (Invitrogen, 65002). Following capture, the newly generated ends were end-repaired and A-tailed using the NEBNext UltraTM II DNA Library Prep Kit for Illumina, and finally ligated to hairpin adaptor 2. After the second adaptor ligation, libraries were prepared by digesting the hairpins on both adapters with USER enzyme (NEB, M5505L) and PCR amplified for 16 cycles using TruSeq index adapters. Sequencing was performed on an Illumina NovaSeq (150 bp paired-end reads).

### Genome alignment

END-seq reads were trimmed of adapters and filtered for reads with low-quality bases using Cutadapt v4.5 with parameters -m 55 -q 25 ^71^. Experimental reads were aligned to the human reference genome (GRCh38) and spike-in reads to *Gallus gallus* (GRCg6a) using bowtie2 v2.5.0 with parameters -N 1 - L 32 ^72^. Processing of aligned reads was performed with samtools v1.17 ^73^, and bedtools v2.31.0 ^74^.

### Spike-In

A set of consensus peaks that occurred in the spike-in reads of all samples was found by taking the overlap of peaks identified with MACS3 v3.0.0b3 using parameters --keep-dup all --nomodel -- nolambda -p 1e-5 ^75^. The normalization factor for each sample was defined as the ratio of the smallest number of reads within consensus peaks in any sample to that sample’s number of reads within the consensus peaks.

### Peak Calling

Significant peaks for each sample were found for the positive strand and negative strand separately, on spike-in-normalized reads with blacklist regions removed^76^. MACS3 v3.0.0b3 was used with parameters --keep-dup all --nomodel --nolambda -p 1e-5, and peaks with fold-change greater than 10 were kept. Peaks were defined as DSBs if the negative-strand peak was within 1000 bp upstream of the positive-strand peak, with the summit of the two peaks between 30 bp and 2500 bp apart.

### Motif Analysis

Recurrent motifs near DSBs were identified with STREME^77^. Fasta files for input to STREME were centered to the taller summit of the DSB, with 1 kb to either side, and the parameter width of 25 was used. Identified motifs were located and scored using FIMO^78^. STREME and FIMO were both used from the MEME Suite v5.5.2 command line software package^79^. A motif was attributed to a DSB if it had a FIMO q-value < 0.05, FIMO score > 0, and occurred within 500 bp of the peak summit. If more than one motif met this criterion, only the the closest motif to the peak was selected.

### Quantitation and statistical analysis

All data presented are biological replicates unless otherwise stated. The statistical tests used, number of replicates, definition of error bars and center definitions are all defined within each figure or figure legend.

## Acknowledgements

We thank Rachel Szilard for critical reading of the manuscript and Jacob Corn for kindly sharing results prior to their publication. We also thank and L. Pelletier, R. Scully for sharing key reagents and Andre Nussenzweig for useful suggestions about END-seq. Work in the DD lab was supported by a grant from the Canadian Institutes for Health Research (CIHR, grant PJT 180438) with additional support from the Reiss Institute for Healthy Aging.

## Author Contributions

Sumin Feng: Conceptualization, Investigation, Writing and Visualization; Lisa Hoeg: Formal Analysis, Data Curation, Visualization; Kaiwen Liu: Investigation, Formal Analysis; Jinfeng Shang: Investigation; Sabrina Roy: Investigation; William Yang: Investigation; Jordan Young: Conceptualization, Supervision; Wei Wu: Validation, Formal Analysis; Dongyi Xu: Supervision, Resources, Funding acquisition; Daniel Durocher: Conceptualization, Supervision, Writing, Funding acquisition.

---

**Figure S1.**
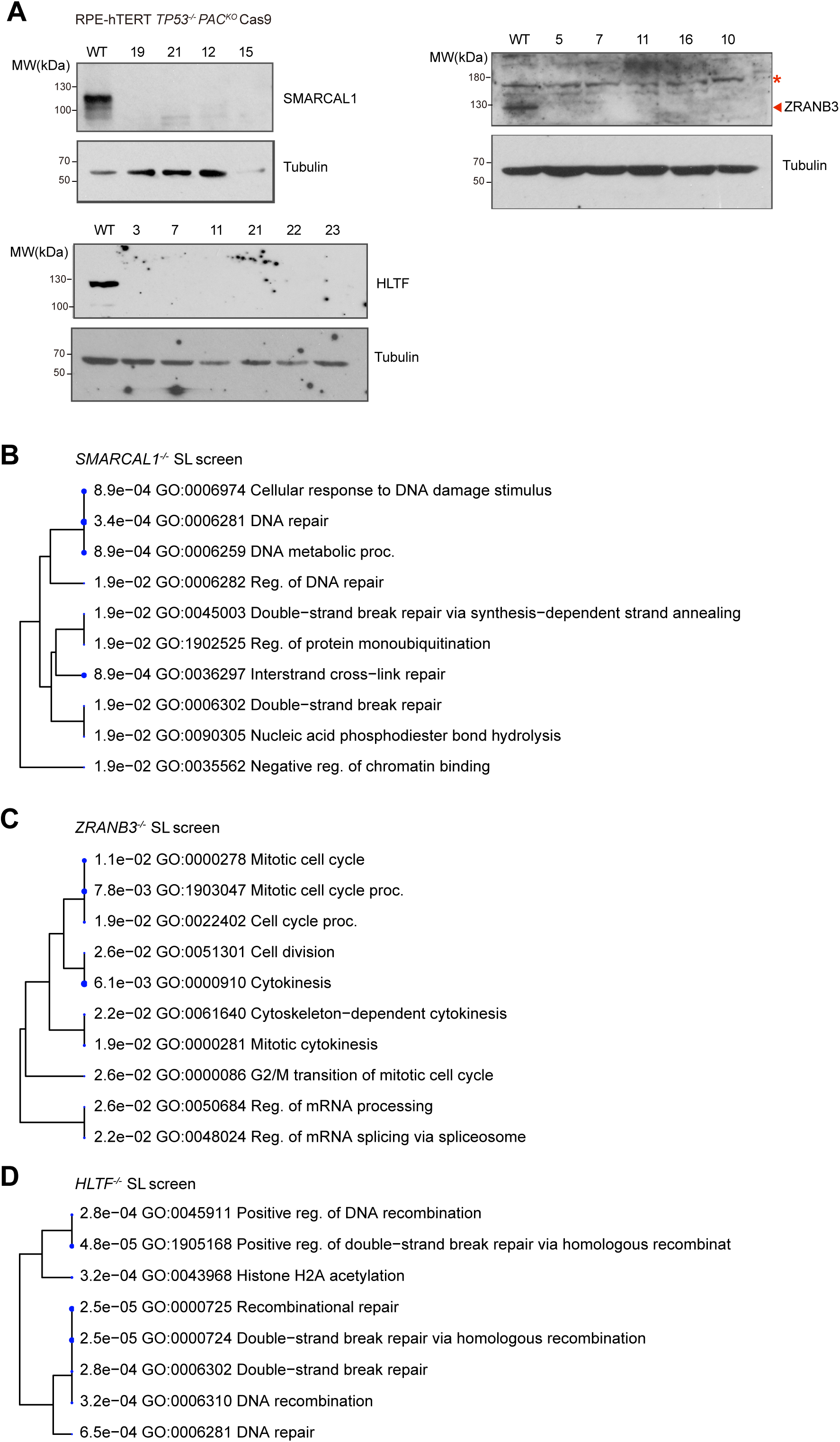
Analysis of knockout cell lines and GO term enrichment in the SNF2-family synthetic lethality screens. (A) Immunoblots examining SMARCAL1 (left), ZRANB3 (right) and HLTF (bottom) levels in RPE1-hTERT *TP53^-/-^PAC^KO^* Cas9 parental (WT) and indicated isogenic gene knockout clones. Tubulin was used as a loading control. Red asterisk indicates non-specific band. Arrowhead indicates specific band. (B-D) GO term enrichment in the *SMARCAL1^-/-^* (B), *ZRANB3^-/-^* (C) and *HLTF^-/-^* (D) screens.

**Figure S2.**
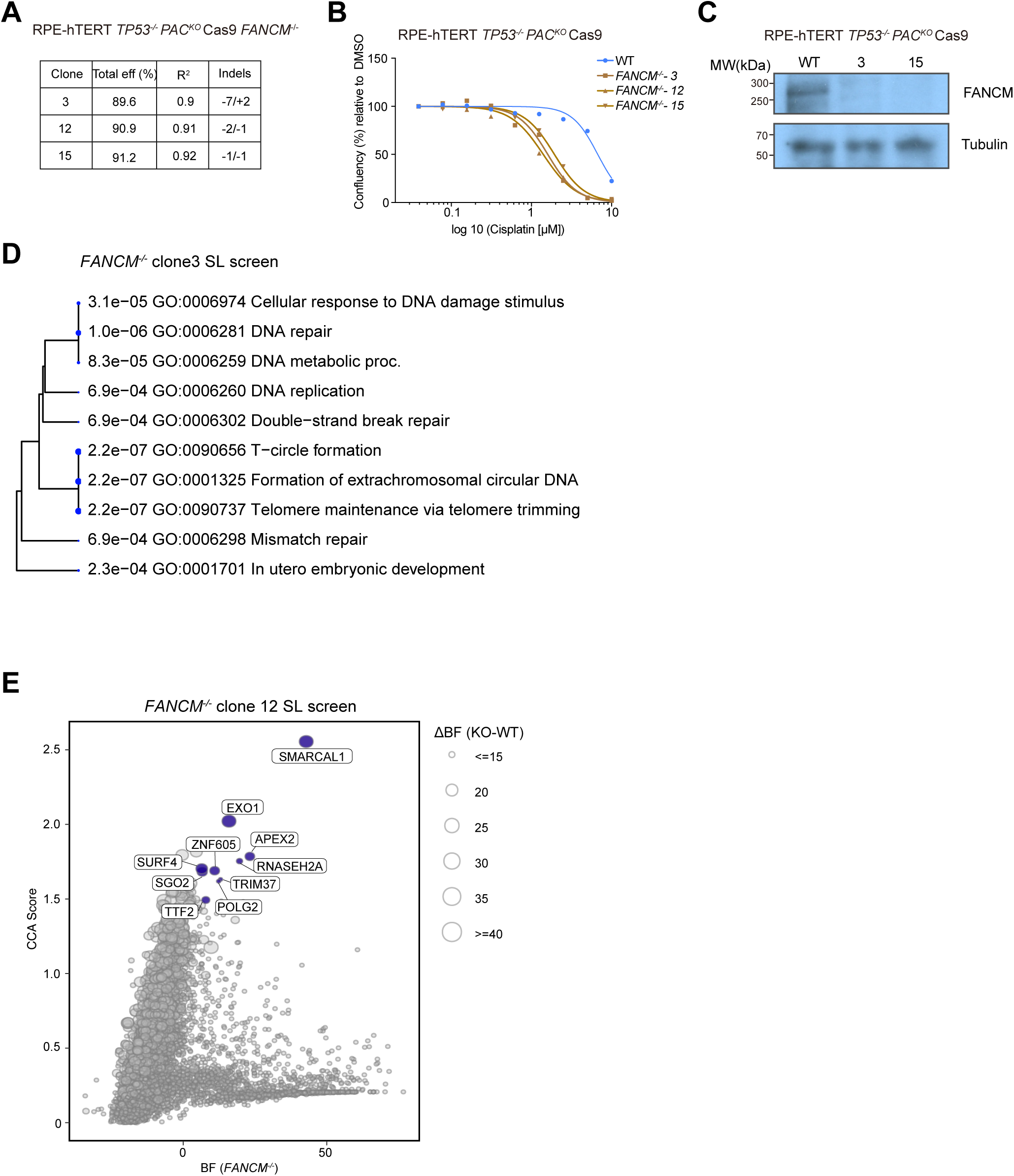
Analysis of *FANCM^-/-^* synthetic lethality screens. (A) *FANCM^-/-^* isogenic clone generation and sanger sequencing screening. Indels is the bps of insertion (+) or deletion (-) on alleles. “/” is used to separate INDELs among different alleles. (B) Cisplatin dose-response assays using confluency as a readout among RPE1-hTERT *TP53^-/-^ PAC^KO^* Cas9 parental (WT) and *FANCM^-/-^*clones (#3, #12 and #15). (C) Immunoblot examining FANCM levels in RPE1-hTERT *TP53^-/-^ PAC^KO^* Cas9 parental (WT) and indicated isogenic gene knockout clones. Tubulin was used as a loading control. (D) As in Figure S1A, but summarizing significant GO terms in *FANCM^-/-^* clone 3 screen. (E) Scatter plot of CRISPR Count Analysis (CCA) scores (y-axis) and Bayes factor (BF) values derived from BAGEL2 (x-axis) for the *FANCM^-/-^* clone 12 isogenic synthetic lethal screen performed in RPE1-hTERT *TP53^-/-^ PAC^KO^* Cas9 cells.

**Figure S3.**
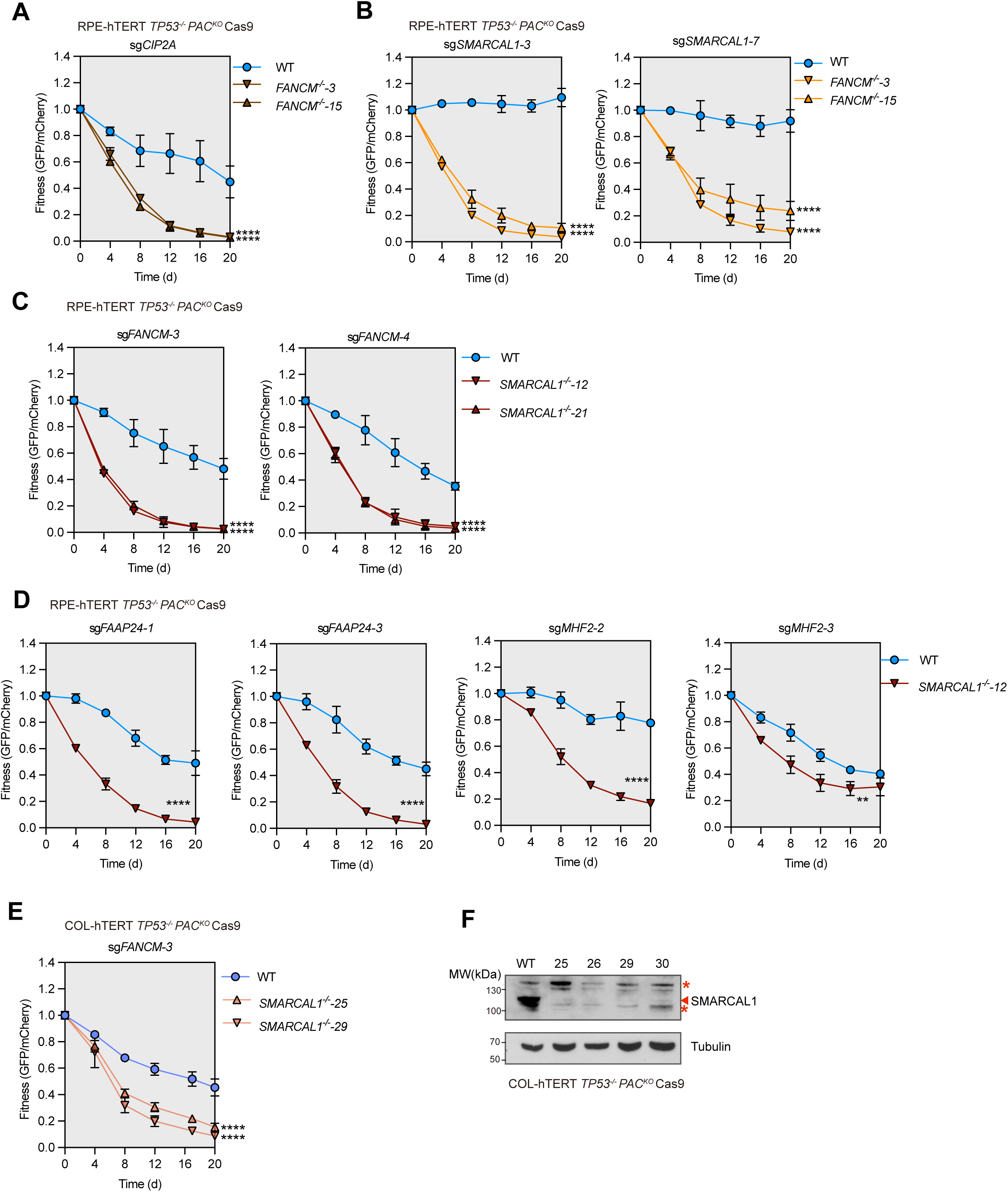
Characterization of the SMARCAL1/FANCM synthetic lethality. (A) Competitive growth assays in RPE-hTERT *TP53^−/−^PAC^KO^* Cas9 cells of the indicated genotypes transduced with a virus expressing an sgRNA targeting *CIP2A*. Data is presented as the mean ± SD (n=3) and was analyzed with a two-way ANOVA test comparing the knockout cell line to the parental line (WT). ****p < 0.0001. (B) As in (A) but using two sgRNAs targeting *SMARCAL1*(#3 and #7). (C) As in (A) but using two sgRNAs targeting *FANCM* (#3 and #4). (D) As in (A) but using two sgRNAs targeting *FAAP24* (#1 and #3) and *MHF2* (#2 and #3), respectively. **p < 0.01, ****p < 0.0001. (E) Competitive growth assays in COL-hTERT *TP53^-/-^ PAC^KO^* Cas9 cells of the indicated genotypes transduced with a virus expressing an sgRNA targeting *FANCM*. Data is presented as the mean ± SD (n=3) and was analyzed with a two-way ANOVA test. ****p < 0.0001. (F) Immunoblot examining SMARCAL1 levels in COL-hTERT *TP53^-/-^PAC^KO^* Cas9 parental (WT) cells and *SMARCAL1* gene knockout clones. Tubulin was used as a loading control. Red asterisk, non-specific band. Arrowhead, specific band.

**Figure S4.**
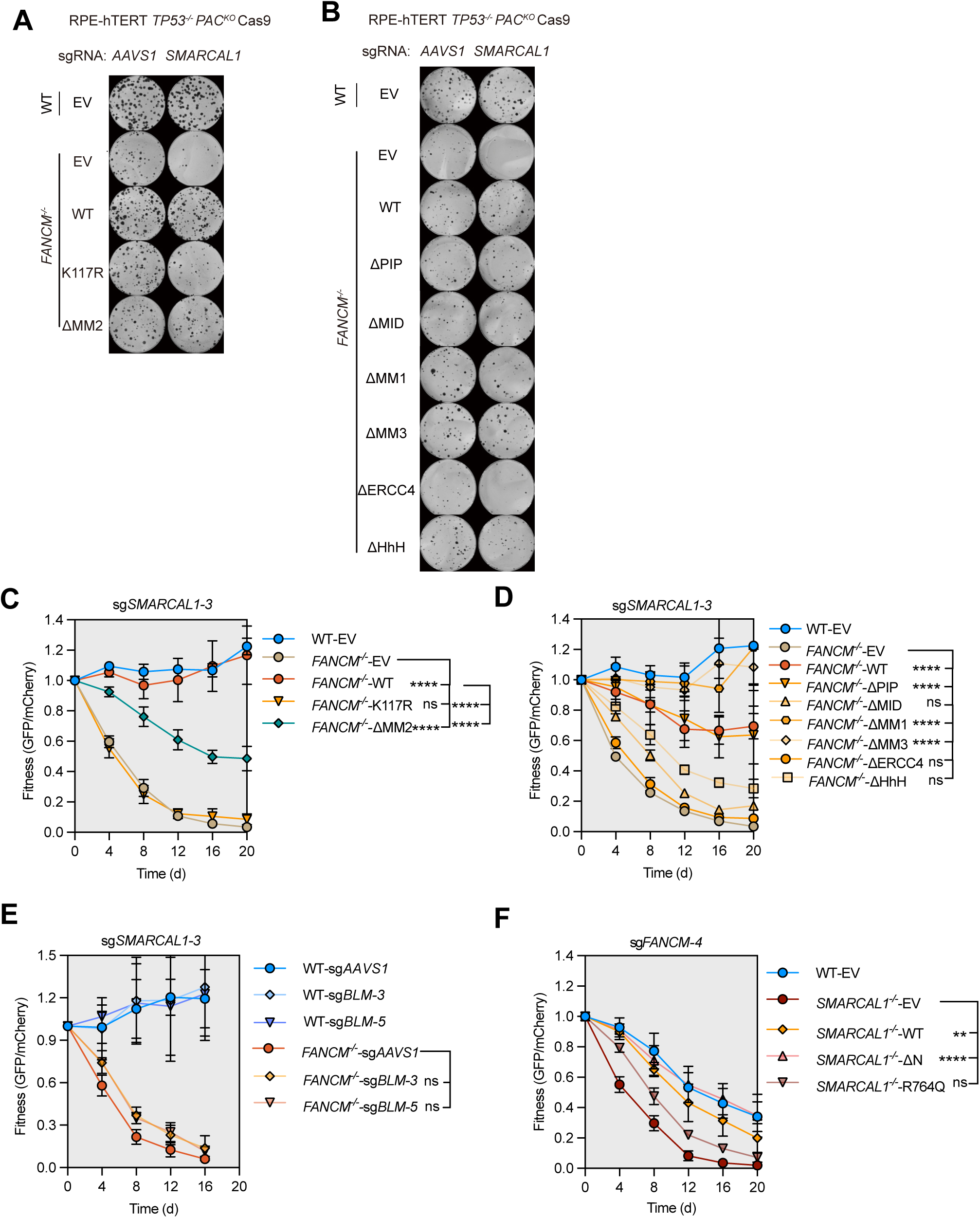
ATPase activity of FANCM or SMARCAL1 is required for cell viability in the absence of the other factor. (A,B) Representative images of clonogenic survival assays in RPE-hTERT *TP53^-/-^PAC^KO^* Cas9 parental (WT) or *FANCM^-/-^* cells expressing the wild type (WT) FANCM, or the indicated variant and transduced with an sgRNA targeting *AAVS1* (control) or *SMARCAL1*. (D) Competitive growth assays of RPE-hTERT *TP53^-/-^ PAC^KO^* Cas9 WT or *FANCM^-/-^* cells with transgenes expressing an empty vector (EV), FANCM WT or its K117R and ΔMM2 mutants and also transduced with an sgRNA targeting *AAVS1* (control) or *SMARCAL1*. Data is presented as the mean ± SD (n=3) and was analyzed with a two-way ANOVA test. ****p < 0.0001, ns, not significant. (E) As in (C) except cells were expressing an empty vector (EV), FANCM WT or its other mutants (ΔPIP, ΔMID, ΔMM1, ΔMM3, ΔERCC4 and ΔHhH). (F) Competitive growth assays of RPE1-hTERT *TP53^-/-^ PAC^KO^* Cas9 WT or *FANCM^-/-^* cells with transgenes expressing and sgRNAs targeting *AAVS1* (control) or *BLM* (#3 and #5). These cells were then transduced with an sgRNA targeting *AAVS1* or *SMARCAL1*. Data is presented as the mean ± SD (n=3), and was analyzed with a two-way ANOVA test. ns, not significant. (G) Competitive growth assays of RPE1-hTERT *TP53^-/-^ PAC^KO^* Cas9 WT or *SMARCAL1^-/-^* cells with transgenes expressing SMARCAL1 WT or mutants (ΔN and R764Q) and also transduced with an sgRNA targeting *AAVS1*(control) or *FANCM*. Data is presented as the mean ± SD (n=3), and was analyzed with a two-way ANOVA test. ****p < 0.0001, **p < 0.01, ns, not significant.

